# Differences in metabolic and liver pathobiology induced by two dietary mouse models of nonalcoholic fatty liver disease

**DOI:** 10.1101/2020.06.05.137174

**Authors:** Hannah Zhang, Mélissa Léveillé, Emilie Courty, Aysim Gunes, Bich Nguyen, Jennifer L. Estall

**Author notes:** Correspondence and reprint requests should be addressed to: Jennifer L. Estall, PhD, Molecular Mechanisms of Diabetes, Institut de Recherches Cliniques de Montreal., 110 avenue des Pins Ouest, Montreal, Quebec, H2W 1R7, Canada. Author Contribution: HZ and JLE designed studies. HZ acquired, analyzed and interpreted data. ML, EC, AG contributed intellectual content. BN scored liver histological samples. HZ and JLE wrote the manuscript. All authors reviewed the manuscript.

## Abstract

Non-alcoholic fatty liver disease (NAFLD) is a growing epidemic associated with key aspects of metabolic disease such as obesity and diabetes. The first stage of NAFLD is characterized by lipid accumulation in hepatocytes, but this can further progress into non-alcoholic steatohepatitis (NASH), fibrosis or cirrhosis, and hepatocellular carcinoma (HCC). A western diet, high in fats, sugars and cholesterol is linked to NAFLD development. Murine models are often used to experimentally study NAFLD, as they can display similar histopathological features as humans; however, there remains debate on which diet-induced model most appropriately and consistently mimics both human disease progression and pathogenesis. In this study, we performed a side-by-side comparison of two popular diet models of murine NAFLD/NASH and associated HCC: a high fat diet supplemented with 30% fructose water (HFHF) and a western diet high in cholesterol (WDHC), comparing them to a common grain-based chow diet (GBD). Mice on both experimental diets developed liver steatosis, while WDHC-fed mice had greater levels of hepatic inflammation and fibrosis than HFHF-fed mice. In contrast, HFHF-fed mice were more obese and developed more severe metabolic syndrome, with less pronounced liver disease. Despite these differences, WDHC-fed and HFHF-fed mice had similar tumour burdens in a model of diet-potentiated liver cancer. Response to diet and resulting phenotypes were generally similar between sexes, albeit delayed in females. Notably, although metabolic and liver disease phenotypes are often thought to progress in parallel, this study shows that modest differences in diet can significantly uncouple glucose homeostasis and liver damage. In conclusion, long-term feeding of either HFHF or WDHC are reliable methods to induce NASH and diet-potentiated liver cancer in mice of both sexes; however, the choice of diet involves a trade-off between severity of metabolic syndrome and liver damage.

## INTRODUCTION

Non-alcoholic fatty liver disease (NAFLD) is a growing epidemic, affecting an estimated one billion people worldwide [1]. Incidence of NAFLD is highly correlated with metabolic diseases, including obesity, diabetes and insulin resistance [1, 2]. The “Western Diet”, characterized by ultra-processed foods rich in saturated fats and refined sugars paired with over-nutrition, is linked to NAFLD initiation and progression [2]. The first stage of NAFLD is simple steatosis and is relatively asymptomatic; however, this can progress to more severe stages of disease over time [3]. Non-alcoholic steatohepatitis (NASH), develops in 30-40% of NAFLD patients and is characterized by liver steatosis in addition to cellular damage and inflammation that can be accompanied by fibrosis and/or cirrhosis [2]. NASH-related hepatocellular carcinoma (HCC) occurs in 4-27% of cirrhotic patient cases, and while this is usually preceded by a cirrhotic stage, HCC can develop in the absence of cirrhosis particularly when coincident with metabolic disease [4].

The genetics of NAFLD are complex; however, data linking the role of different dietary components in NAFLD pathobiology is strong [5]. Specific dietary components influence lipid accumulation in the liver and NAFLD progression. While glucose is the major circulating carbohydrate source for energy, there has been a large increase in the consumption of fructose via processed foods and sugary drinks [6]. Consumption of a high-fructose diet increases hepatic lipid content by enhancing lipogenesis and also promotes reactive oxygen species (ROS) formation, contributing to liver steatosis and NAFLD progression [7]. Diets high in fat, when consumed in conjunction with high carbohydrates, also contribute to hepatic lipid accumulation [7]. Saturated fats in particular promote oxidative stress and hepatocyte death [8]. Dietary cholesterol is linked to NASH progression, as increased free cholesterol in hepatocytes increases ROS production [9], activates Kupffer cells to release inflammatory cytokines and promotes fibrosis through hepatic stellate cell (HSC) activation [10]. High-cholesterol feeding also leads to aberrant hepatic gene expression, specifically in the pathways of calcium, insulin, cell adhesion, axon guidance and metabolism, and promotes NASH-related HCC in mice [11]. Changes that accompany liver inflammation and fibrosis can lead to DNA damage and cellular reprogramming, and can create a pro-tumourigenic environment [12].

Pathogenic mechanisms of NASH are not fully understood and treatments other than diet modification and exercise are not well-established [13]. Our understanding of human NAFLD aetiology and pathobiology is complicated due to disease heterogeneity and difficulty of early diagnosis. Murine models are a useful model to study NAFLD and NASH, as they can display similar hepatic histopathologic characteristics to humans [14].

Although studies in mice must be carefully interpreted for translatability, these models have provided crucial insights into the evolution of NAFLD/NASH progression [15]. However, the choice of mouse model is a significant topic for debate in the field. It is generally accepted that an ideal mouse model will mimic both the pathophysiology and histopathology of human disease progression [16]. In this regard, there are 2 main criteria we believe murine models of NAFLD/NASH should encompass in addition to liver steatosis, including: (1) liver damage (including both inflammation and fibrosis), and (2) manifestation of metabolic disease, such as obesity and/or insulin resistance. Currently, there is no consensus in the field on a mouse model that meets all the above factors. Scientists often face a trade-off between modeling the metabolic characteristics of NAFLD versus producing a NASH liver phenotype, and are discouraged by the time required to induce liver phenotypes in mice.

A diet-based model that mimics nutrient type and relative quantities consumed by humans is generally favoured over genetic or chemically-induced NASH models. Many murine diets used for NASH do not cause obesity or insulin resistance (eg. methionine choline-deficient diet) [17]. High-fat and/or sucrose diets traditionally used to induce metabolic syndrome and NAFLD in mice can be highly variable, and the degree of liver inflammation and fibrosis is generally low [15]. The Gubra amylin NASH (GAN) diet is an emerging model of NASH high in saturated fats, sucrose and cholesterol. Male C57Bl/6J mice fed the GAN diet have severe liver histopathology similar to NASH [18], but to our knowledge, it is not known whether diets of this type induce levels of obesity and insulin resistance commonly associated with high-fat/high-sucrose feeding in mice or humans. Human data also suggests separate etiologies of NAFLD/NASH in males and females [19]; yet direct comparisons of diet effect on metabolic and liver phenotypes between sexes is not available, limiting mechanistic insight and translatability of the animal model.

This study aims to physiologically compare two commonly used diet-induced mouse models of obesity and metabolic syndrome with NAFLD in both sexes. The high-fat high-fructose (HFHF) model is a high-fat/high sucrose diet supplemented with 30% fructose (w/v) in the drinking water. This model is relevant to human consumption patterns given current consumption of a significant proportion of sugar in liquid form (e.g. soft drinks or juices) [6]. Metabolic and liver profiles of mice on this diet were compared side-by-side with littermate mice on a “Western Diet” high in fat, fructose and cholesterol (WDHC), very similar to the GAN diet [18], and a popular grain-based rodent diet (GBD) used commonly for colony maintenance in animal facilities. Our study shows that the high-fat/high sugar diets both induce obesity and NAFLD, but there were important differences in the extent of obesity, development of metabolic syndrome, and severity of liver pathology that were diet- and sex-dependent.

## MATERIALS AND METHODS

### NASH Cohort Diets and Mice

All mice were wild-type on an inbred C57BL/6N genetic background. Animals were housed on a 12-hour light/12-hour dark cycle and fed the respective diets *ad libitum*. The grain-based diet (GBD) (Teklad Global 18% Protein Rodent Diet) was obtained from Teklad Diets, Envigo (Huntingdon, UK), and consists of 24 kcal% protein, 58 kcal% carbohydrates and 18 kcal% fat. A pelleted high fat/high sucrose diet (D12451i, ResearchDiets, USA), composed of 20 kcal% protein, 35 kcal% carbohydrates and 45 kcal% fat, was supplemented with 30% D-fructose (BioShop, Burlington, ON) drinking water to create the high-fat/high-fructose (HFHF) diet. The western diet high in cholesterol (WDHC) (D17010103i, ResearchDiets, USA) was composed of 20 kcal% protein, 40 kcal% carbohydrates and 40 kcal% fat, with 2% w/w cholesterol. *Table S1* compares the macronutrient composition of each diet side-by-side. 18 male mice (n=4 on GBD; n=7 on HFHF; n=7 on WDHC) and 22 female mice (n=6 on GBD, n=8 on HFHF; n=8 on WDHC) were started on their respective diets at 6-8 weeks of age. All weeks are denoted as weeks on the diet unless otherwise indicated. Body mass was taken weekly. At the end of the experimental protocol, mice were sacrificed following a 3-6 hour fast. Cardiac puncture was performed for blood collection. Livers and pancreata were isolated and weighed. Samples were fixed in formalin for histology or frozen in liquid nitrogen and stored at -80°C until extraction of proteins, lipids and RNA.

### HCC Mouse Cohort

Wild-type inbred C57BL/6N genetic background mice were housed in the same fashion as the NASH cohort mice described above. Mice were injected with 25 mg of diethylnitrosamine (DEN) per kg body mass at 2 weeks of age. Mice were fed *ad libitum* either the HFHF diet or the WDHC diet as described above. 14 males (n=6 on HFHF, n=8 on WDHC) and 14 females (n=7 on HFHF and n=7 on WDHC). Body mass was taken weekly, and mice were sacrificed after 24 weeks of feeding following a 3-hour fast. Samples were processed in a similar manner as the NASH cohort.

### Blood Collection

A minimum concentration of 3.2 µL per 600 µL blood of Aprotinin from bovine lung (Sigma, MA, USA) was added to all blood samples (except those taken during the glucose tolerance test) to prevent protein degradation. Blood collected during the oral glucose tolerance test (OGTT) was collected in *Microvette ® 100µL K3 EDTA* (Sarstedt, Germany). Blood was incubated at 4°C for 90 minutes, followed by an 8-minute centrifugation at 4000 rpm at 4°C. The top layer of serum was collected and stored at -80°C until analysis.

### Circulating ketone, insulin and lipid quantification

Serum ketone β-Hydroxybutyrate was measured using a colorimetric kit (Cayman Chemical, MI, USA). Aminoalanine transferease (ALT), was measured using the *Liquid ALT (SGPT) Reagent Set* (Pointe Scientific Inc., MI, USA) as per manufactures instructions.

Insulin levels were measured by enzyme-linked immunosorbent assay (ELISA) using the *Mouse Ultrasensitive Insulin ELISA* (ALPCO, NH, USA) on serum samples taken at 0 and 15 minutes during the OGTT, and at the study endpoint (after 31 weeks on the diets).

Lipids were measured directly in serum or following extraction from liver tissue using 2:1 chloroform:methanol followed by 60% butanol, 40% solution of 2:1 Triton X-114:methanol. Total acyl-glycerol content was determined using the *Total Triglyceride Reagent Kit* (Sigma, MA, USA). Free glycerol values were subtracted from total glycerol concentration to obtain acyl-glycerol content. Non-esterified fatty acid (NEFA) levels were assessed using the *HR Series NEFA-HR(2)* kit (Fujifilm Wako Chemicals U.S.A., VA, USA). Cholesterol levels were quantified using the Cholesterol Reagent Set (Pointe Scientific Inc, MI, USA), with a cholesterol standard obtained from the *Cholesterol Assay Kit* (Abcam, Cambridge, UK).

### Magnetic Resonance Imaging (MRI)

Live animals were weighed prior to MRI using the *minispec mq7*.*5 NMR analyzer* (Bruker, MA, USA) to obtain fat and lean mass data at 12, 24 and 31 weeks following initiation of diet protocols.

### Respirometry

Mice were housed individually in metabolic cages of the *Promethion Metabolic Measurement System* (SableSystems International, NV, USA) between 4 and 10 weeks on their respective diets for a total of 7 days. Only the final three full light/dark cycles of data were used for analyses. VO_2_ (mL), VCO_2_ (mL), energy expenditure (kcal), food and water intake (grams and mL, respectively) and pedometer movement (meters) were continuously measured by the system. Respiratory exchange ratio (RER) is the ratio of VCO_2_/VO_2._

### Oral Glucose and Insulin Tolerance Tests

Oral glucose tolerance tests (OGTT) were performed after 16 weeks of diet feeding. Mice were fasted overnight for 16 hours. 1.5 g glucose/kg body weight was administered by oral gavage. Blood glucose levels were measured in tail blood using a *Freestyle Lite* glucometer (Abbott, IL, USA) at 0, 15, 30, 45, 60, 90 and 120 minutes following glucose administration. Tail blood was collected at 0 and 15 minutes post-glucose administration for insulin content analysis. Insulin Tolerance Tests (ITT) were performed after 22 weeks on the diets. Mice were fasted for 5 hours prior to intraperitoneal injection with insulin *Humulin R* (Eli Lilly, IN, USA) at 0.7 IU/kg body weight or 0.5 IU/kg body weight for males and females, respectively. Blood glucose levels were taken from tail vein using a *Freestyle Lite* glucometer (Abbott, IL, USA) at 0, 15, 30, 45, 60, 90 and 120 minutes post-injection.

### Quantitative Polymerase Chain Reaction (qPCR)

Total RNA was extracted from the left medial lobe of the liver using *TRIzol*^*®*^ *Reagent* (Ambion, Thermo Fisher Scientific, MA, USA) according to the manufacturers protocol. Samples were homogenized and 1 µg of RNA was treated with DNaseI prior to cDNA synthesis using the *High Capacity Reverse Transcription Kit* (Applied Biosystems, Carlsbad, CA). A Reverse Transcriptase negative control was included to control for genomic DNA contamination. *SensiFAST*^*™*^ *SYBR® Lo-ROX Kit* (Bioline, TN, USA) was used to amplify cDNA using primers listed in *Table S2*, and quantified using a *ViiA7 Real-Time PCR System* (ThermoFisher, MA, USA). ΔΔCt threshold cycle method of normalization was used, with genes normalized to hypoxanthine-guanine phosphoribosyltransferase (HPRT) and expressed relative to levels in grain-based diet fed mice.

### Histology and Scoring

The left lateral lobe of the liver and pancreata were fixed in formalin and embedded in paraffin. Hematoxylin and eosin (H&E) and Sirius red staining were performed on tissues and slides were visualized using transmission brightfield microscopy. A pathologist assessed simple steatosis, hepatocyte ballooning, lobular inflammation and fibrosis in blinded sections, according to the NAFLD Activity Score-Clinical Research Network (NAS-CRN) [20]. Details in scoring are found in Figure 1 and 2.

**Figure 1.**
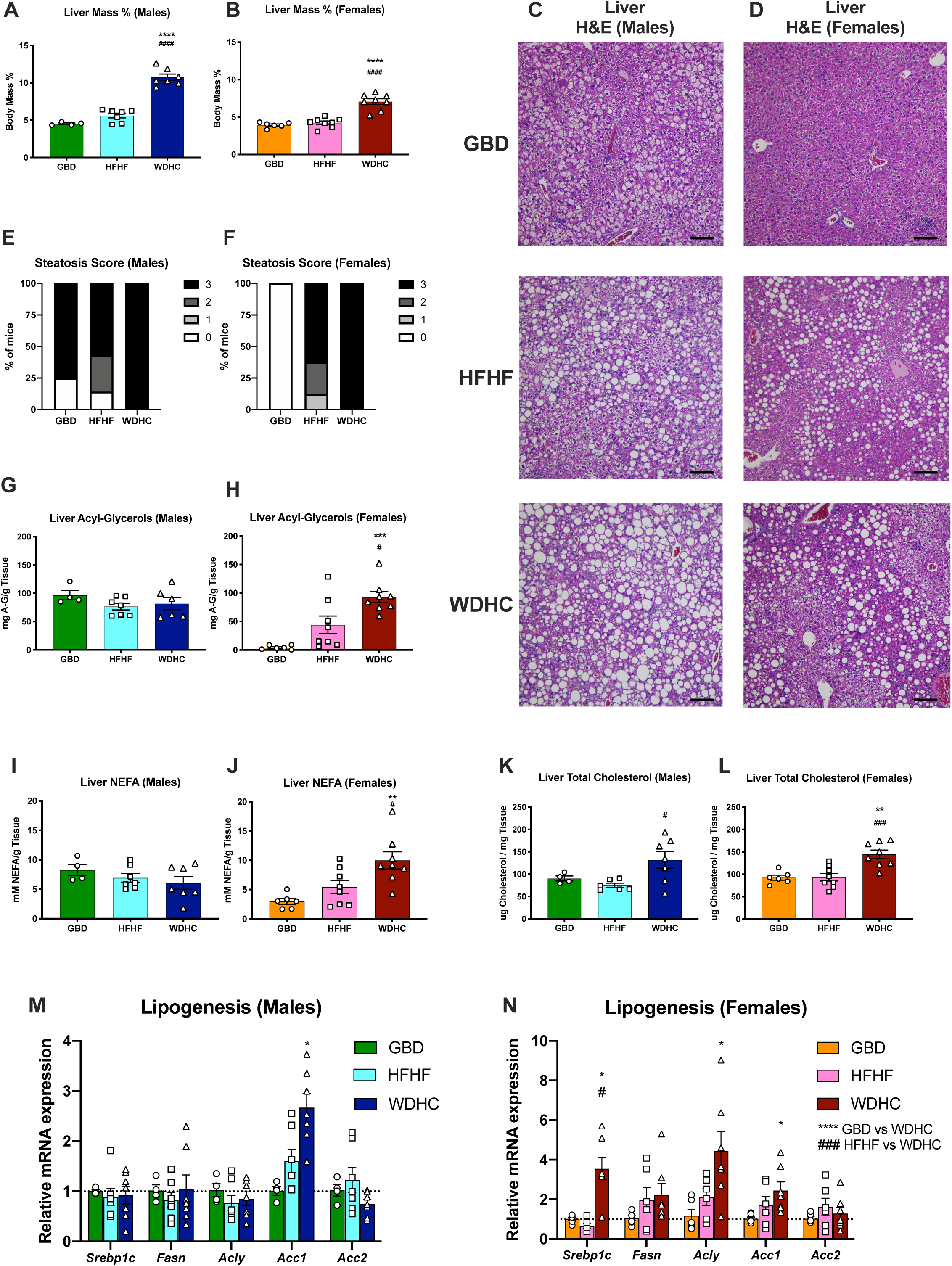
WDHC feeding causes hepatomegaly and more pronounced induction of lipogenic gene expression compared to HFHF diet. (A-B) Liver mass as a percentage of body mass at sacrifice (31 weeks of feeding). (C-D) Hematoxylin & Eosin (H&E) staining of liver tissue at 31 weeks of feeding (bars represent 100µm). (E-F) Grading of liver steatosis from 0-3 (0: <5% steatosis, none; 1: 5-33%, mild; 2: 34-66%, moderate; 3: >67%, marked). (G-H) Liver acyl-glycerol content (I-J); liver non-esterified fatty acid (NEFA) levels; and (K-L) liver total cholesterol levels at sacrifice. (M-N) Hepatic lipogenic gene expression measured by qPCR. Data are expressed as mean ± SEM (n=4-8) and representative of two individual cohorts. Results of Two-way ANOVA and t-tests are reported in panels. ^*^p≤0.05, ^**^p ≤ 0.01, ^***^p ≤ 0.001, ^****^p ≤ 0.0001 compared to GBD; ^#^p≤0.05, ^###^p ≤ 0.001, ^####^p≤0.0001 comparing WDHC to HFHF.

**Figure 2.**
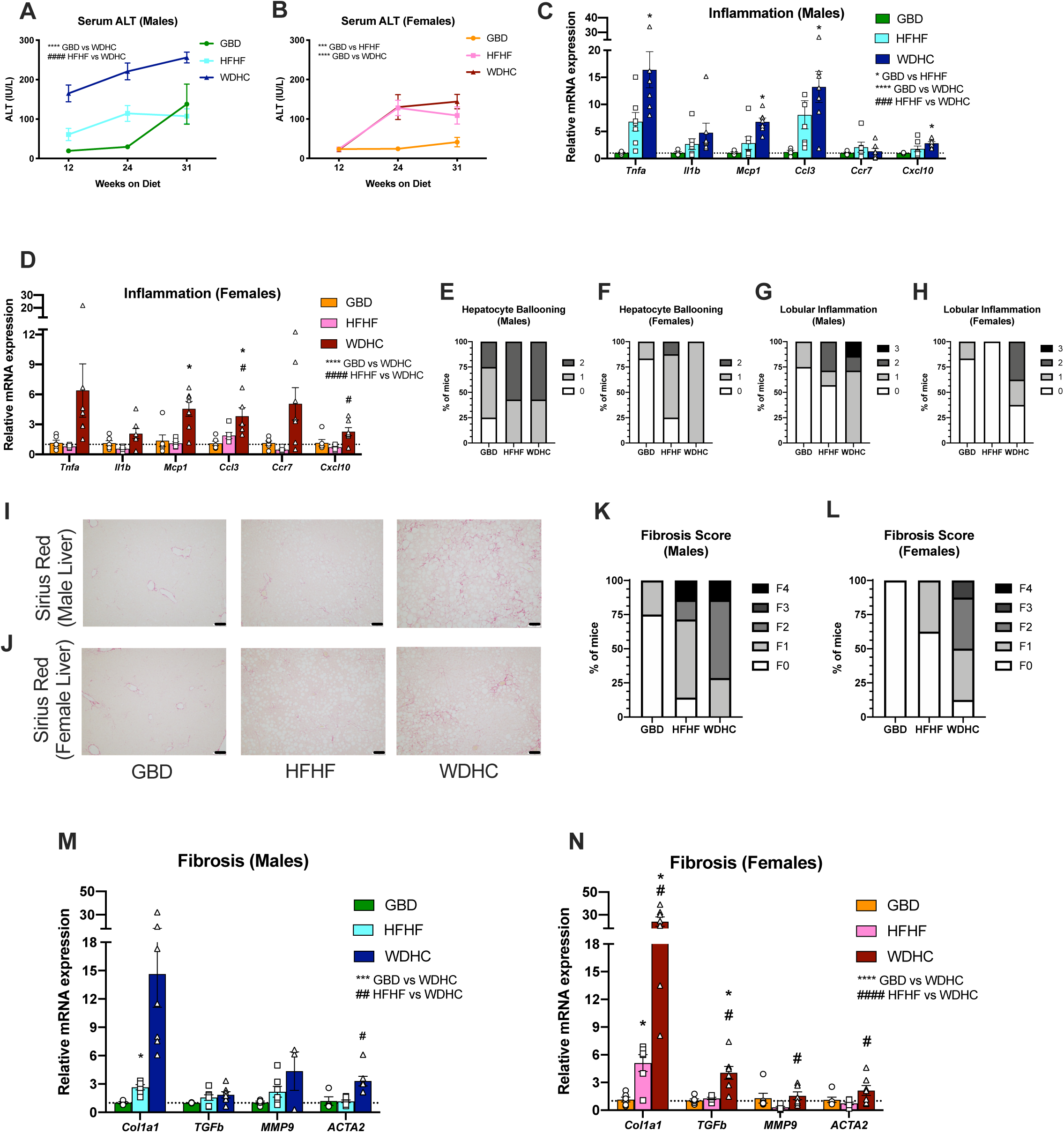
WDHC-fed mice have increased liver damage. (A-B) Serum alanine aminotransferase (ALT) levels. (C-D) Hepatic inflammatory gene expression measured by qPCR. (E-F) Scoring of hepatocyte ballooning from 0-2 (0: none; 1: few balloon cells; 2: many cells/prominent ballooning) and (G-H) scoring of lobular inflammation, based on number of inflammatory foci per 20X field (0: no foci; 1: <2 foci; 2: 2-4 foci; 3: >4 foci). (I-J) Sirius Red staining of liver tissue taken at sacrifice (31 weeks of feeding), bars represent 100µm. (K-L) Stage of fibrosis using NAS-CRN scoring (F0: no fibrosis; F1: perisinusoidal zone 3 or portal fibrosis; F2: perisinusoidal and periportal fibrosis without bridging; F3: bridging fibrosis; F4: cirrhosis). (M-N) Fibrotic gene expression in mouse livers measured with qPCR. Data are expressed as mean ± SEM (n=4-8) and representative of two individual cohorts. Results of Two-way ANOVA and t-tests are reported in panels. ^*^p ≤ 0.05, ^**^p ≤ 0.01, ^***^p ≤ 0.001, ^****^p ≤ 0.0001 compared to GBD; ^#^p ≤ 0.05, ^##^p ≤ 0.01; ^###^p ≤ 0.001; ^####^p ≤ 0.0001 comparing WDHC to HFHF.

### Statistics

All data are shown as mean ± standard error of the mean (SEM). GraphPad Prism (GraphPad Software, CA, USA) was used for all statistical analyses. Unmatched 2-way analysis of variance (ANOVA) was used for all time-course parameters, with Sidak’s multiple comparison test with a significance cut-off of p<0.05. For qPCR data, genes were separated into pathways prior to analysis by 2-way ANOVA, followed by multiple t-tests correcting for multiple comparisons using the Holm-Sidak method. Unmatched one-way ANOVA test was used for comparison of parameters at a single-timepoint in the NASH Cohort, using the Tukey test to correct for multiple comparisons. T-tests were used to compare diet effects at a single timepoint in the HCC cohort of mice. Outliers, determined by Grubb’s test (GraphPad Prism), were excluded from analysis.

## RESULTS

### HFHF- and WDHC-fed livers were both steatotic, but showed differences in size, lipids, and lipogenic gene induction

NAFLD is characterized by the presence of significant steatosis encompassing >5% of hepatocytes [20]. After 31 weeks of diet feeding, both male and female mice fed the Western diet high in cholesterol (WDHC) had disproportionately larger livers than HFHF-fed mice when normalized to body mass (*Fig. 1A, B*). Hematoxylin and Eosin (H&E) staining of liver tissue for both males and females showed high levels of steatosis for both HFHF and WDHC (*Fig. 1C, D*). WDHC-fed mice of both sexes had higher steatosis scores compared to their GBD- and HFHF-fed counterparts (*Fig. 1E, F*). In males, liver acyl-glycerol and non-esterified fatty acid (NEFA) levels after 31 weeks of feeding were not significantly different between any of the diets (*Fig. 1G, I*). WDHC-fed female mice had higher liver acyl-glycerol and NEFA concentrations than female mice fed the GBD or HFHF diet (*Fig. 1H, J*). Both males and females fed WDHC had higher hepatic cholesterol levels compared to HFHF-fed mice (*Fig 1K, L*). Increased lipogenic gene expression is reported for both mouse models of NALFD and human NASH [21, 22]. In male mice, the only difference noted in hepatic lipogenic gene expression was an increase in Acetyl-CoA carboxylase 1 (*Acc1*) expression for WDHC-fed mice (*Fig. 1M*). Similar increases in *Acc1* were seen in WDHC-fed females, but this was accompanied by higher hepatic expression of Sterol regulatory element-binding transcription factor 1 (*Srebp1c)* and ATP citrate lyase (*Acly)* compared to mice fed GBD or HFHF (*Fig. 1N*). Together, these results indicate that both HFHF and WDHC diets cause steatosis, but there were appreciable differences in the extent of lipid accumulation and effect on lipogenic pathways at the time of sacrifice.

### WDHC-fed mice had increased inflammation and liver damage

To determine if the diets recapitulated liver damage commonly associated with NASH, we characterized hepatic injury, inflammation and fibrosis. Serum ALT levels are a nonspecific marker for liver damage [23]. In male mice, WDHC feeding induced higher levels of circulating ALT compared to the other diets throughout the course of the study, suggesting early induction of liver damage (*Fig. 2A*). HFHF-fed male mice also had higher ALT levels at 12 and 24 weeks of feeding, although GBD-fed mice had levels similar to HFHF-fed males by the end of the study (*Fig. 2A*). While 12 weeks on the diet was not sufficient to induce serum ALT levels in females on the different diets, after 24 and 31 weeks HFHF- and WDHC-fed females both had elevated ALT levels compared to GBD-fed females (*Fig. 2B*). ALT levels for both males and females fed the HFHF diet appeared to peak around 24 weeks.

Liver inflammation is also a key hallmark of NASH [8]. Hepatic expression of inflammatory cytokines commonly associated with NASH were upregulated in WDHC-fed mice of both sexes (*Fig. 2C, D*), as were markers of inflammatory cell populations including monocytes, B-cells and T-cells (*Fig. S1*). Hepatic inflammatory markers in HFHF-fed mice were increased to a lesser extent than WDHC-fed mice (*Fig. 2C, D*), and there were sex differences in the type and level of immune cell response (*Fig. S1*). Hepatocyte ballooning and lobular inflammation scores were both higher in male and female mice fed WDHC compared to GBD and HFHF (*Fig. 2E-H*). Fibrosis is an indication of more progressed liver disease and is a key prognostic factor in human NASH [24]. Sirius Red staining of liver tissue revealed increased levels of collagen deposition only in WDHC-fed mice in both males and females (*Fig. 2I, J)*. Fibrosis scores were higher in HFHF-fed mice compared to GBD-fed mice, and higher in mice fed WDHC compared to both other diets (*Fig. K, L*). We also observed increased fibrotic gene expression in the WDHC-fed mouse livers of both sexes (*Fig. 2M, N*). Thus, these results suggest that WDHC-fed mice had significant levels of inflammation and liver fibrosis at the time of sacrifice.

### Mice chronically fed HFHF diet were larger, and females more obese, than mice fed WDHC

NAFLD and NASH are strongly associated with obesity [25]. Both male and female mice fed HFHF diet were heavier than WDHC- and GBD-fed mice (*Fig. 3A, B*). Female WDHC-fed mice weighed significantly more than GBD-fed females (*Fig. 3B*), while WDHC-fed males gained weight early and plateaued, eventually weighing significantly less than the other two diets (*Fig. 3A*). The effect of each diet on weight gain was exaggerated in females following chronic feeding (>15 weeks) compared to differences observed in males. MRI analysis of body composition after 12 weeks showed a slight trend for increased fat mass in male WDHC-fed mice compared to males fed GBD (*Fig. S2A*). Fat mass values in HFHF-fed males were highly variable. Unexpectedly, males fed GBD had high amounts of body fat. Female mice showed early increases in fat mass on both HFHF and WDHC diets (*Fig. S2B*). Of note, absolute lean mass (g) was also increased for males fed HFHF (*Fig. S2C*) and females fed either diet (*Fig. S2D*) compared to their GBD-fed counterparts at 12 weeks. After 31 weeks of feeding, GBD-fed mice were notably obese (30% fat mass) and there were no differences in fat mass for male mice in any group (*Fig. 3C*). Females fed HFHF and WDHC had significantly higher fat mass percentages compared to GBD-fed mice, with HFHF-fed females being the most obese (*Fig. 3D*). At this later time point, both HFHF- and WDHC-fed males (*Fig. 3E*), and HFHF-fed females (*Fig. 3F*), had higher absolute lean mass (g) compared to GBD-fed mice. As higher lean mass may indicate larger mice overall, body length was measured from the tip of the snout to the base of the tail. Females on HFHF were significantly longer than GBD-fed females, with similar trends seen for WDHC and for both diets in males (*Fig. 3G, H*).

**Figure 3.**
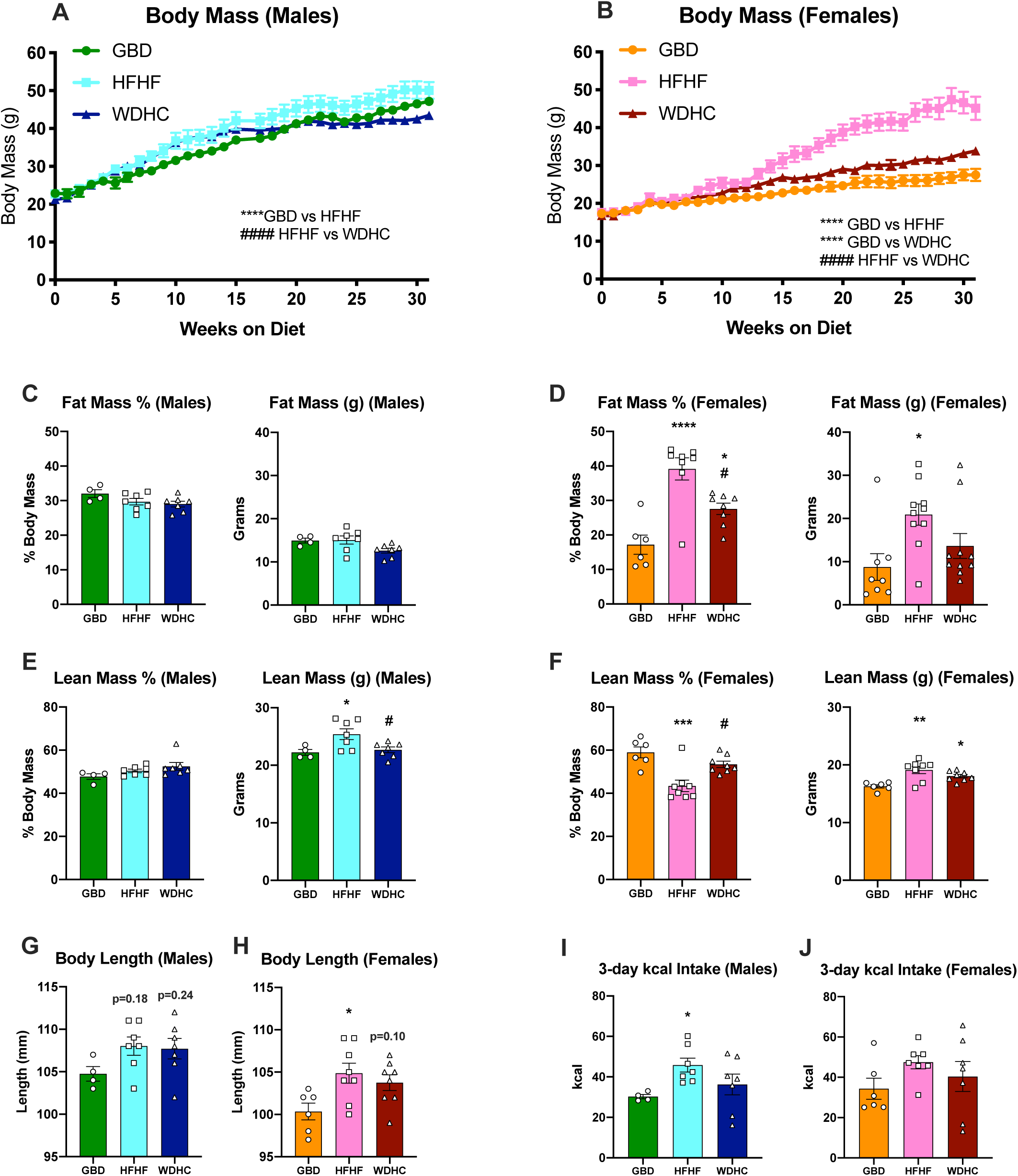
HFHF-fed mice are larger, and females more obese, than WDHC-fed mice. (A-B) Body mass measured weekly. (C-D) Fat mass normalized to total body mass and absolute fat mass by MRI. (E-F) Normalized lean mass and absolute lean mass taken by MRI. (G-H) Body length measured after 14 weeks of diet feeding. (I-J) Total caloric intake measured over 3 days in metabolic cages, calculated from recorded food and water intake. Data are expressed as mean ± SEM (n=4-8) and representative of two individual cohorts. Results of Two-way ANOVA and One-way ANOVA are reported in panels. ^*^p≤0.05, ^**^p ≤ 0.01, ^***^p ≤ 0.001, ^****^p ≤ 0.0001 compared to GBD; ^#^p ≤ 0.05, ^####^p ≤ 0.0001 comparing WDHC to HFHF.

To determine whether differences in body mass were due to variation in energy intake and expenditure, we used metabolic cages to assess the metabolic activity of the mice after 4-10 weeks feeding, before weights had significantly diverged (*Fig*.*3A, B*). There were no significant differences in 3-day total solid food or water intake observed between mice on the different diets, even when normalized to body mass (*Fig. S3A-H*). The 3-day total caloric intake (including water plus food) of mice on the HFHF diet was higher in males and trended higher in females, due to added calories from water consumption (*Fig. 3I, J*). Total energy expenditure and total pedometer movement over 3 days was not different between diets in either group (*Fig. S3I-L*). Overall, HFHF and WDHC diets significantly increased weight of both sexes. Increases in weight were due to changes in both fat mass and lean mass, while HFHF-feeding led to significantly more fat mass in female mice compared to WDHC. Differences in body composition and weight were not due to changes in feeding behaviour, movement, or total energy expenditure; however, increased intake of carbohydrate (fructose) via the drinking water was likely a significant contributing factor. Taken together, the sex of the mice, in conjunction with diet type, significantly influenced differences in weight gain and body composition.

### WDHC-fed mice have increased fat metabolism

Circulating lipid levels are determined by the coordinated processes of fat storage and catabolism, and high levels of serum lipids can indicate inefficient fat storage or increased lipolysis. In both males and female serum acyl-glycerol levels in WDHC-fed mice were lower compared to GBD-fed mice. A similar trend was seen for the HFHF-fed mice, although not statistically significant (*Fig. 4A, B*). No differences in serum NEFA levels were observed between the diets in either sex, although WDHC-fed males trended lower (*Fig. 4C, D*).

**Figure 4.**
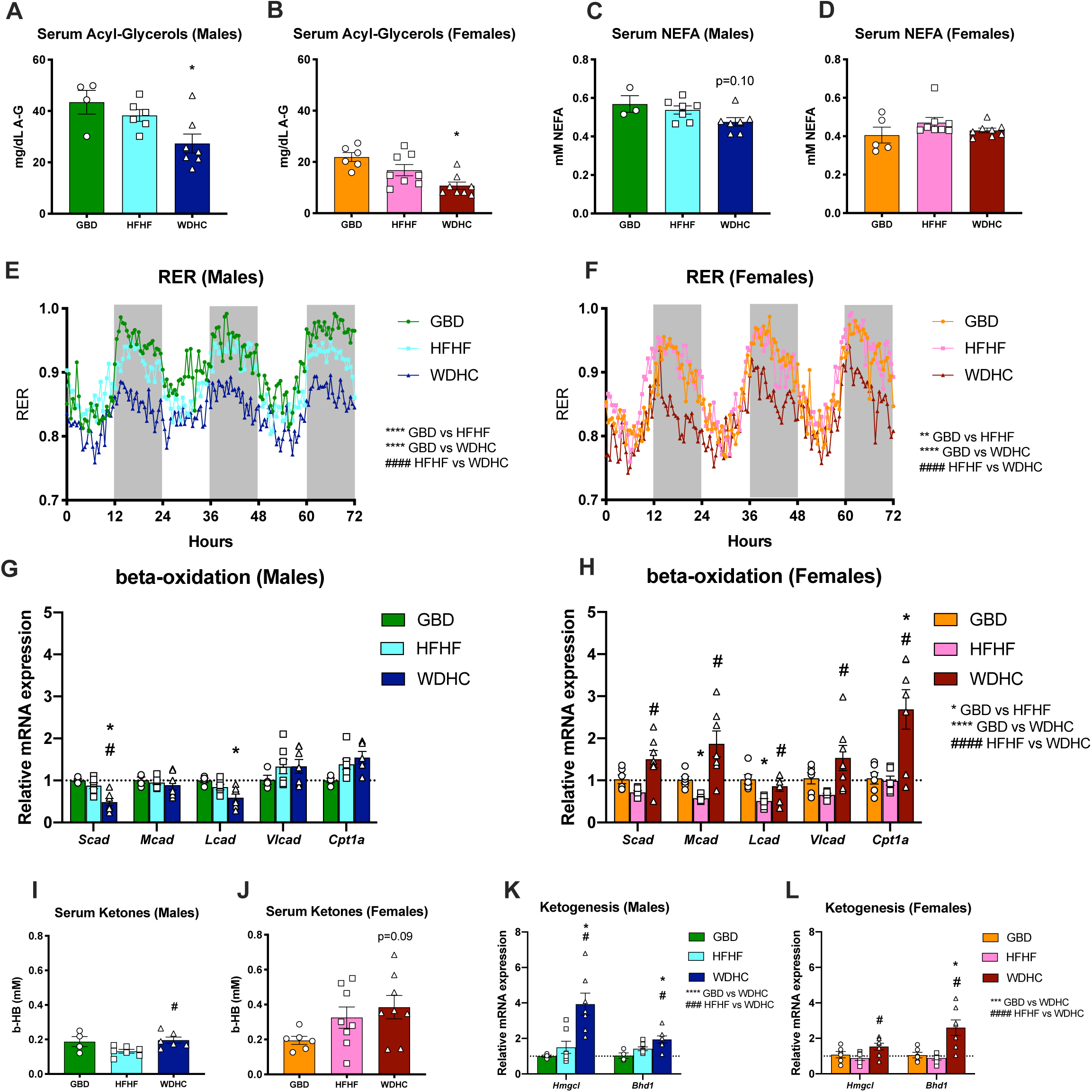
WDHC-fed mice have increased fat metabolism compared to HFHF-fed mice. (A-B) Serum acyl-glycerol levels measured at sacrifice (31 weeks of feeding). (C-D) Serum NEFA levels quantified at 31 weeks of feeding. (E-F) Respiratory exchange ratio (RER) measured by metabolic cages over 3 days. (G-H) Liver beta-oxidation gene expression measured by qPCR. (I-J) Serum ketone beta-hydroxybutyrate (b-HB) levels measured at 31 weeks of feeding. (K-L) Liver ketogenic gene expression measured by qPCR. Data are expressed as mean ± SEM (n=4-8) and representative of two individual cohorts. Results of Two-way ANOVA and t-tests are reported in panels. ^*^p≤0.05, ^**^p ≤ 0.01, ^***^p ≤ 0.001, ^****^p ≤ 0.0001 compared to GBD; ^#^p ≤ 0.05, ^####^p ≤ 0.0001 comparing WDHC to HFHF.

Although the HFHF and WDHC diets had similar total fat and carbohydrate content (*Table S1*), respiratory exchange ratios (RERs) were lower in WDHC-fed mice in both sexes, suggesting increased utilization of lipid as an energy source over carbohydrates compared to GBD- and HFHF-fed mice, particularly in the dark cycles (*Fig. 4E, F*). We then sought to know if the low RER of WDHC-fed mice was due to increased β-oxidation in liver. Female WDHC-fed mice were the only group to have increased expression of multiple genes within the β-oxidation pathway including Short-, Medium, Long, and Very-long chain acyl-CoA dehydrogenase (*Scad, Mcad, Lcad, Vlcad*) and Carnitine Palmitoyltransferase 1A (*Cpt1a*) compared to GBD after 31 weeks on the diet (*Fig. 4G, H*). In contrast, hepatic mRNA levels in WDHC-fed males were mostly unchanged except for a reduction in *Scad* and *Lcad* expression, despite similar decreases in RER between sexes. β-oxidation genes were also generally reduced in livers of female mice fed HFHF, but not males (*Fig. 4G, H*). Ketone bodies, such as β-hydroxybutyrate (β-HB), are produced using acetyl-CoA derived from fatty acid oxidation. WDHC-fed mice of both sexes had a trend towards higher circulating β-HB levels compared to HFHF-fed mice, although not statistically significant in females due to the high variability (*Fig. 4I, J*). In both males and females, there was an upregulation of ketogenic genes HMG CoA lyase (*Hmgcl*) and β-HB dehydrogenase (*Bhd1*) in WDHC-fed mouse livers compared to GBD- and HFHF-fed mouse livers (*Fig. 4K, L*). Increased hepatic β-oxidation and ketogenesis in WDHC-fed female mice may contribute to their lower RER. Increases in beta-oxidation pathways were not seen in male livers; however, mRNA expression changes were measured in liver tissues many weeks following metabolic cage analysis, limiting our ability to conclude on direct changes in liver metabolism occurring at the time of RER measurement. However, decreased RER early on and increased markers of ketogenesis in both male and female mice on WDHC at the end of the experiment suggest a shift toward a more fasting-like or ketotic state of lipid metabolism with this diet, involving lipid precursors coming from liver, as well as other tissues (i.e. adipose).

### Glucose intolerance and liver pathology did not correlate well for any diet

Glucose intolerance and insulin resistance are clinically relevant features in NAFLD, as they are associated with each other [26]. Oral Glucose tolerance tests (OGTTs) and insulin tolerance tests (ITTs) were performed in mice on the respective diets to determine the efficiency of glucose clearance. As expected, HFHF-fed males and females were less glucose tolerant than GBD-fed mice (*Fig. 5A, D*). In stark contrast, WDHC-fed male mice were significantly more glucose tolerant than mice fed either GBD or HFHF diets. Their increased glucose clearance may be explained by increased glucose-stimulated insulin secretion (*Fig. 5B*) combined with increased insulin sensitivity (*Fig. 5C*). In females, WDHC diet had no effect on glucose tolerance (*Fig. 5D*) or secreted insulin levels (*Fig. 5E*), but significantly increased insulin sensitivity (*Fig. 5F*). Of note, while male mice on the GBD had a poor response to insulin comparable to mice on the HFHF diet, this is in line their similar body mass and composition at that time (*Fig. S2*). Generally, WDHC-fed mice of both sexes appear metabolically healthier than the HFHF-fed mice, with better glucose tolerance and insulin sensitivity.

**Figure 5.**
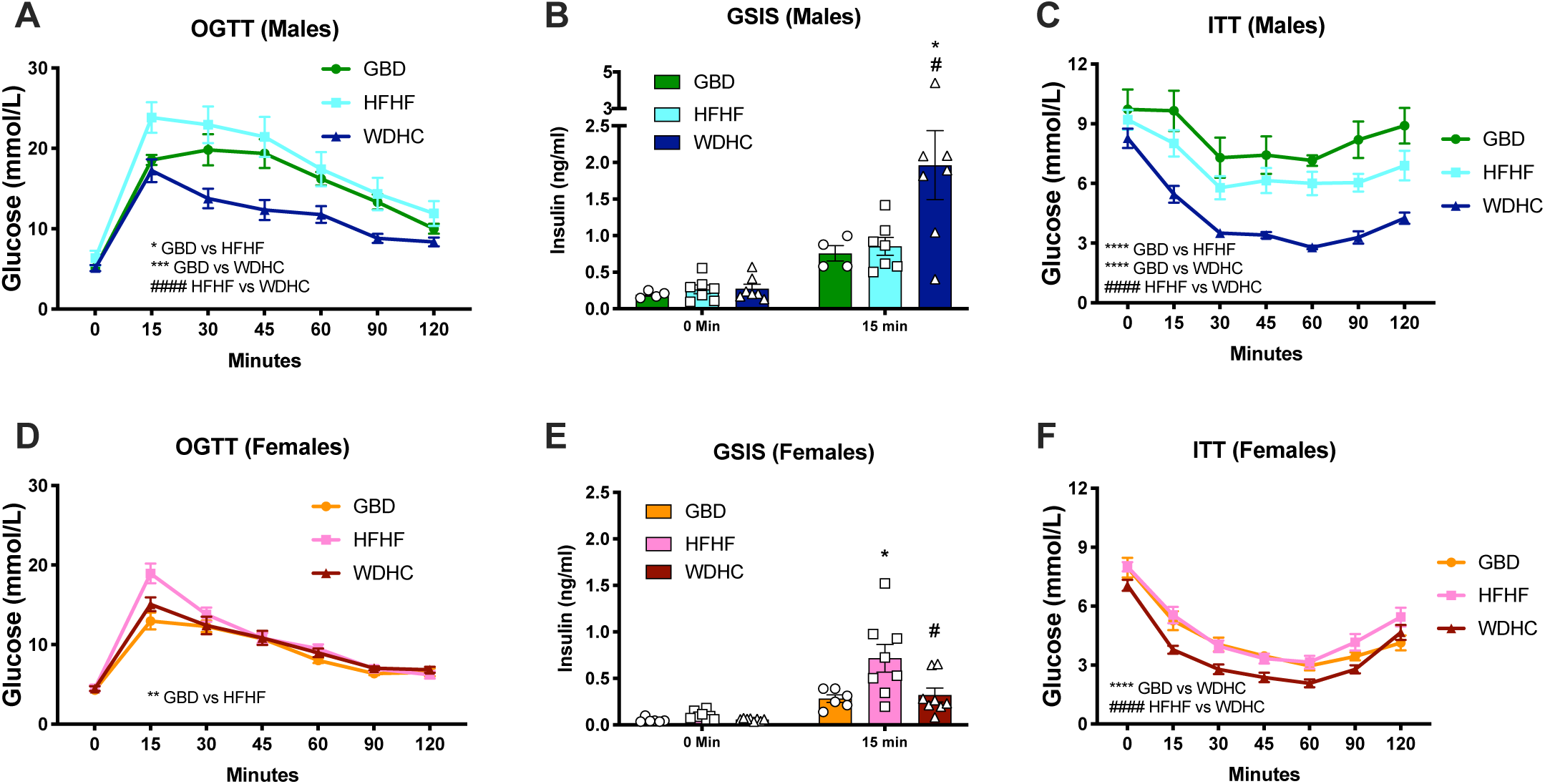
WDHC and HFHF diets have opposing effects on glucose homeostasis. (A, D) Oral glucose tolerance test (OGTT) performed at 16 weeks of feeding, and (B, E) the associated glucose stimulated insulin secretion (GSIS) measured by ELISA. (C, F) Insulin tolerance test (ITT) performed at 22 weeks of feeding. Data are expressed as mean ± SEM (n=4-8) and representative of two individual cohorts. Results of Two-way ANOVA are reported in panels. ^*^p≤0.05, ^**^p ≤ 0.01, ^***^p ≤ 0.001, ^****^p ≤ 0.0001 compared to GBD; ^#^p ≤ 0.05, ^####^p ≤ 0.0001 comparing WDHC to HFHF.

### HFHF and WDHC induce NASH-related HCC

In a state of damage, liver cells induce regenerative pathways to repair damaged tissue [8], which can create a pro-tumorigenic environment [12]. In our NASH cohort of mice, we assessed hepatic expression of genes associated with proliferation and apoptosis to determine if this could account for the increased liver mass (*Fig. 1A, B)* and high levels of liver damage (*Fig. 2*) observed in WDHC-fed mice. In male mice, there was no overall difference in proliferative gene expression in livers, but female mice fed WDHC had increased hepatic expression of Proliferating cell nuclear antigen (*Pcna*) and Myc-related translation/localization regulatory factor (*Myc*) (*Fig. 6A, B*). Hepatic expression of anti-apoptotic genes B-cell lymphoma 2 (*Bcl2*), B-cell lymphoma-extra large (*BclXL)*, and *Survivin* (*Birc5*) were increased in WDHC-fed mouse livers of both sexes compared to GBD-fed mouse livers (*Fig. 6C, D*). Expression of DNA damage-inducible transcript 3 (*Ddit3*, commonly known as *Chop*) (a pro-apoptotic transcription factor and marker of increased endoplasmic reticulum stress) was increased in WDHC-fed mice compared to GBD-fed mice, but not HFHF-fed mice (*Fig. 6E, F*). There were no significant increases in mRNA for pro-apoptotic markers Bcl2 associated agonist of cell death (*Bad*) or X Protein (*Bax*) for either diet. These data suggest increased cell proliferation and decreased apoptosis in WDHC livers, correlating with increased liver weight.

**Figure 6.**
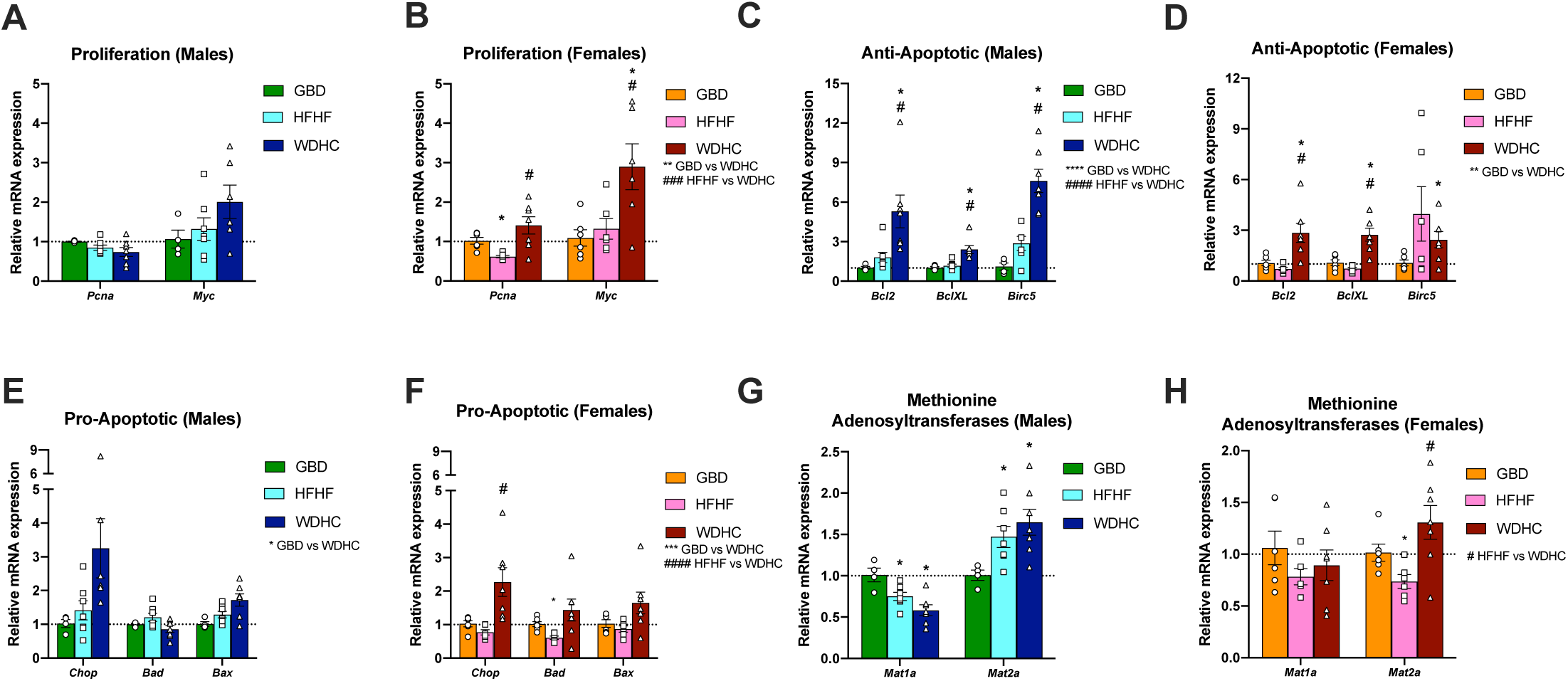
WDHC-fed mice displayed signs of increased liver turnover. qPCR of mRNA levels for gene representing (A-B) proliferation; (C-F) apoptosis; and (G-H) S-adenosylmethionine synthesis in mouse livers at sacrifice. Data are expressed as mean ± SEM (n=4-8) and representative of two individual cohorts. Results of Two-way ANOVA and t-tests are reported in panels. ^*^p≤0.05, ^**^p ≤ 0.01, ^***^p ≤ 0.001 compared to GBD; ^#^p ≤ 0.05, ^####^p ≤ 0.0001 comparing WDHC to HFHF.

Given the evidence of dysregulated hepatocyte growth and survival, we next investigated whether the expression of methionine adenosyltransferases (*Mat*s) were altered. *Mat1a* is a marker of mature liver cells, mainly expressed in differentiated hepatocytes and bile duct cells, commonly downregulated in HCC [27, 28]. *Mat2a* is primarily found in fetal liver, but its expression can be induced in adult livers during periods of rapid liver growth, proliferation and dedifferentiation, such as HCC [28]. In male mice fed either the HFHF- or WDHC diets, there was a decrease in hepatic *Mat1a* and an increase in *Mat2a* (*Fig. 6G*). *Mat1a* expression was not altered by the diets in female mice. Similar to males, WDHC feeding increased *Mat2a* expression in female mice (*Fig. 6H*), while HFHF diet reduced *Mat2a* expression in contrast to males. Overall, these results imply that the WDHC diet caused increased liver turnover.

Increased liver turnover could create an environment more conducive to tumour development. To investigate if the WDHC diet provided a more pro-tumourigenic environment, we employed a model of chemical carcinogen-induced HCC in conjunction with the NAFLD-promoting diets. Diethylnitrosamine (DEN) is a DNA alkylating agent biotransformed and activated by CYP450 enzymes in liver. It causes liver tumour development in mice after 4-6 months when given concurrently with a potentiating agent (i.e. phenobarbital, streptozocin or high caloric diets) [29]. Mice on GBD were not included, as they generally do not develop liver tumours in this model. Similar to the NASH cohort, HFHF-fed mice receiving DEN were significantly heavier than their WDHC-fed counterparts at the end of the study (*Fig. 7A, B*) and WDHC-fed mice had disproportionately larger livers than HFHF-fed mice (*Fig. 7C, D*), regardless of sex. There were no significant differences in fat mass and lean mass percentages at the end of the experiment (*Fig. 7E, F*); however, similar to the trend seen in our NASH cohort (*Fig. S2*), fat mass in WDHC-fed males was increased early in the diet time course (*Fig. S5A, B*). Trends in liver lipids in this cancer cohort (*Fig. S5*) paralleled those observed in our NASH cohort (*Fig. 1*). There was no significant difference in liver acyl-glycerol content or steatosis between the two diets at the end of the study (*Fig. S5E-H*).

**Figure 7.**
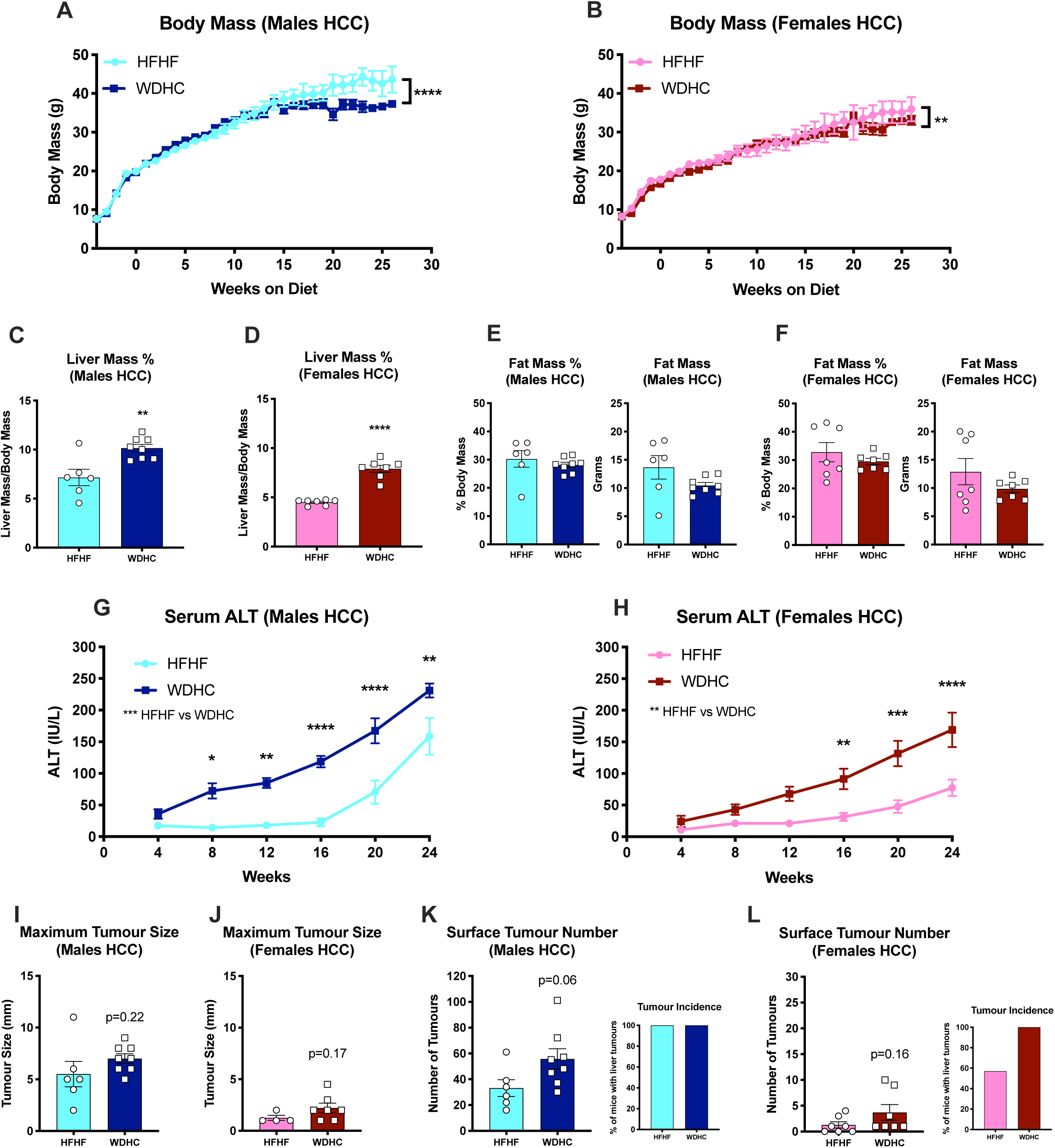
Both HFHF and WDHC diets potentiate liver cancer development. (A-B) Body mass was taken weekly beginning at the time of DEN injection. (C-D) Liver mass at sacrifice normalized to total body mass. (E-F) Fat mass normalized to body mass and absolute fat mass taken with MRI at 24 weeks on the diets; (G-H) Serum ALT levels measured every 4 weeks. (I-J) Maximum liver tumour size taken at sacrifice. (K-L) Surface tumour number counted at sacrifice with tumour incidence expressed as percentage of mice with liver tumours. Data is expressed as mean ± SEM. Data are expressed as mean ± SEM (n=4-8) and representative of two individual cohorts. Results of Two-way ANOVA and t-tests are reported in panels. ^*^p≤0.05, ^**^p ≤ 0.01, ^***^p ≤ 0.001, ^****^p ≤ 0.0001.

In line with increased liver damage and hepatic inflammation, circulating ALT was significantly higher in WDHC-fed mice of both sexes compared to HFHF-fed mice, starting its rise in serum much earlier (*Fig. 7G, H*). WDHC-fed mouse livers had higher inflammatory gene expression in females, although no difference was seen in males (*Fig. S6*). Interestingly, although the diets caused very different phenotypes in terms of glucose homeostasis and liver damage, tumour number and size were similar in both groups. Maximum tumour size (*Fig. 7I, J*) and surface tumour number (*Fig. 7K, L*) trended higher in WDHC-fed mice of both sexes, although the difference was not statistically significant. Importantly, there was 100% incidence of tumours in females on the WDHC diet compared to 57% on the HFHF diet (*Fig. 7K, L*), suggesting that this diet regime could help overcome the poor efficacy often faced when modeling HCC in female mice. Taken together, the HFHF and WDHC diets provoked murine liver tumour development to a similar extent following exposure to DEN, with increased liver damage correlating with a trend in increased tumor load following WDHC feeding.

## DISCUSSION

A diet-induced mouse model of NAFLD/NASH that mimics both human disease pathogenesis and progression would increase translatability to human physiology. We performed a side-by-side comparison of the liver and metabolic phenotypes of mice of both sexes on two commonly used diet-induced models of NAFLD/NASH and assessed their relative efficacy to potentiate HCC. While mice fed WDHC had increased liver damage and fibrosis, HFHF-fed mice had more severe metabolic phenotypes including obesity and insulin resistance. We also showed that while sex seemed to play a role in the timing of disease progression, both sexes developed NASH and diet-potentiated liver cancer following chronic feeding of a diet high in fat, fructose and cholesterol.

Although liver and metabolic phenotypes in the context of NAFLD/NASH are frequently assumed to be concurrent and codependent, this study demonstrated that they can be uncoupled depending on diet composition. This finding is in line with recent data showing that NAFLD/NASH and other metabolic disease phenotypes can be separated [17]. Despite similar fructose consumption between mice fed HFHF and WDHC, a large portion of the fructose consumed by HFHF-fed mice was in liquid form. Liquid fructose consumption can confer increased risk of metabolic syndrome, manifesting in insulin resistance and obesity, compared to solid sugar consumption [30, 31]. This is consistent with the HFHF-fed mice having more advanced metabolic syndrome compared to WDHC-fed mice. The lack of obesity in WDHC-fed mice could be explained by increased catabolism of non-hepatic lipids, such as adipose tissue stores. Consistent with this, RER data, a measurement of whole-body energy utilization, suggests that mice fed WDHC relied more on lipid as a fuel source compared to HFHF-fed or GBD-fed mice. Recent data alternatively suggests fructose in liquid versus solid form differentially alter the gut microbiota and liver toxicity [32, 33], which could lead to differences in whole body lipid catabolism and storage downstream of the different diets. It is important to consider that metabolic cage data, including food/water intake and RER, was collected very early in the protocol, while our analysis of liver histology and gene expression was performed at the end of the study. Thus, it is possible we did not capture important changes in hepatic metabolic pathways occurring at the energy expenditure data was collected.

Our results are in line with the growing evidence that hepatic cholesterol accumulation potentiates liver disease [9], as WDHC-fed mice had increased levels of serum ALT levels, inflammation and fibrosis compared to HFHF-fed mice. Interestingly, there is a distinct population of NASH patients who are not obese and have less pronounced insulin resistance compared to obese NASH patients [34, 35], similar our WDHC-fed mice. WDHC-fed mice had an early increase in ALT, which is interesting considering that lean NAFLD is also associated with younger age on disease onset [36]. Furthermore, increased cholesterol consumption has been observed in patients with non-obese NASH compared to subjects with obese NASH [35]. Therefore, the HFHF diet appears to be a comprehensive model of obesity-related NAFLD/NASH patients, including criteria of liver steatosis, hepatocyte damage, obesity and insulin resistance, while WDHC may represent a good model of non-obese NASH. This hypothesis would benefit from further studies to compare the WDHC diet-induced murine model directly to pathobiology of patients with non-obese NASH, as has been done with previously for obesity-related NASH.

Our data are consistent previous studies emphasizing the need for long feeding protocols to allow progression of NAFLD to NASH in mice. Diet-induced animal model of NAFLD (DIAMOND) mice are obese, insulin resistant and develop liver fibrosis when fed a diet similar to our HFHF model for 24-52 weeks; however, their phenotype relies heavily on genetic background [37]. Our WDHC diet is similar to the GAN diet, both modeled on the now discontinued Amylin Mouse Liver NASH (AMLN) diet. These diets all combine high-fat with high fructose and cholesterol (2% w/w), with key characteristics of NASH starting to appear after 12 weeks on the diet [37, 38]. The inclusion of trans-fat in the original AMLN diet further potentiates hepatic fat storage, ALT appearance and insulin resistance [39]; however, following the Food and Drug Administration’s (FDA) ban on trans-fat addition to food products, this dietary component has become difficult to obtain commercially [40]. While trans-fat is proven to potentiate liver disease, some argue that addition of trans-fat to mouse chow is now less representative of current human diet patterns given their recent decrease in trans-fat consumption [7]. The GAN, WDHC and HFHF diets, and specifically inclusion of fructose or sucrose in the drinking water [31], arguably better mimic the make-up of current human diets compared to previous models. Yet, like in humans, these diet protocols all require long periods of time (greater than 24 weeks of feeding in mice) to cause significant NASH.

Shorter times to disease development are often preferred for preclinical studies NAFLD/NASH studies, to minimize cost and maximize output. Models such as methionine- and/or choline-deficient diets or the Stelic Animal Model (STAM) diet are frequently used to rapidly induce NASH in mice; however, mice under these protocols do not develop significant obesity or insulin resistance, with some experiencing weight loss and increased insulin sensitivity [17, 41, 42]. In contrast, human NAFLD and NASH usually progress over decades, with age being a main risk factor of disease severity [43]. The longer experimental period of 31 weeks feeding in mice provided sufficient time to observe significant liver fibrosis, increases (and plateaus) in circulating ALT, concurrent with increases in fat mass and insulin resistance. In particular, our study showed that with longer feeding periods, female mice also acquired significant liver disease, including NASH-related HCC. Our study provides approximate time windows for the development of metabolic and liver phenotypes in both sexes, and ultimately shows, NAFLD/NASH pathology is possible to model in both sexes using these diets.

Epidemiological and *in silico* data suggest that NAFLD progresses through different mechanisms between sexes [44], although these have not been extensively studied *in vivo* [45]. Estrogen signaling protects females from both metabolic syndrome and liver damage [46]. Female mice typically have decreased levels of estrogen when they are between 39-52 weeks old [47]. Consistent with this, we saw no elevation in circulating ALT in female mice at the early timepoint of 12 weeks on any diet. However, at 24 and 31 weeks, we saw a divergence in ALT levels, with both test diets elevating levels. This is consistent with females having significant liver inflammation and fibrosis by the end of our study, which is not usually observed in females after short-term feeding (when they are included). While this may imply that females simply take longer to acquire liver disease compared to males, it is also possible that females have a separate etiology of liver disease. Our study designed to define times lines and characteristics of liver disease development in male and female mice should facilitate the design of future studies to delineate the important similarities and differences in NASH development between sexes.

Dietary-driven mouse models of NASH-related HCC have similar drawbacks to diet-induced NASH mouse models, often not mirroring human etiology and metabolic features, or having low incidence of tumour development [15]. Our HCC cancer model showed that the combination of an early carcinogen with either HFHF or WDHC feeding are effective models of NASH-related HCC. Although not statistically significant, mice fed WDHC had modestly larger and more numerous tumours than their HFHF-fed counterparts. Importantly, females fed WDHC in this cancer model had 100% incidence of tumours, which may have been a factor limiting the use of female mice in liver cancer studies in the past. The WDHC-feeding model may help to overcome this hurdle and is encouraging for researchers who wish to study pathogenic mechanisms of NASH-related HCC development in mice that are relevant to all humans.

There are a few important caveats and limitations exist to our study. Most notably, males fed our GBD (standard animal facility chow) acquired significant steatosis and insulin resistance compared to data from similar studies using purified control diets [39]. While our original intention was to use a commonly used control chow diet (i.e. use of purified control diets is rare), there was surprisingly little difference in fat mass and insulin sensitivity in male mice fed the GBD chow when compared to HFHF-fed mice at 28-30 weeks of age. This result could be a consequence of aging and not the dietary components; however, WDHC-fed mice of the same age were very sensitive to insulin. It is possible that the GBD used in our study, being relatively higher in fat content (18%) than purified control diets (10%) and containing unspecified sources of carbohydrate, is sufficiently obesogenic on its own. It is worthy to note that despite this caveat, aside from liver phenotypes related to obesity (e.g. lipid levels), liver damage in GBD-fed male mice was less than in HFHF- and WDHC-fed mice, and female mice on the same GBD responded as expected. This observation brings up an important consideration when choosing a chow control diet, as matched purified control diet may provide better predictability of response and facilitate data interpretation.

In summary, the HFHF model may be the superior model for the changes in energy homeostasis typically associated with obesity-related NAFLD and NASH. In contrast, the WDHC diet could be a valuable tool to study NASH-related HCC, as well as non-obese NASH development. This study provides detailed information intended to aid researchers in the choice of mouse model best representing the characteristics of the human population of interest. While there may be no perfect diet to model human NAFLD/NASH in mice, our results provide insights to the vast difference in phenotypes that can be seen with varying dietary components, times of feeding and responses between sexes.

## Supplementary Materials

**Supplementary Table S1.** Composition of the different diets used in this study. GBD= Grain-Based Diet; HFHF=High-Fat High-Fructose; WDHC=Western Diet High in Cholesterol *Values are calculated based on the average daily consumption during the last 3 days of the metabolic cages experiments **Assumes lard is 39% saturated fat and soybean oil is 16% saturated fat

| Component | GBD | HFHF | WDHC |
| --- | --- | --- | --- |
| Constituents | <u>Teklad 2918</u> | D12451i + 30% Fructose Water | D09100103 + Regular Water |
| Total Protein (kcal%) | 24 | 17.15 | 20 |
| Total Carbohydrates (kcal%) | 58 | 44.27 | 40 |
| Total Fat (kcal%) | 18 | 38.58 | 40 |
| Fructose (gm%) | ~0 | 24.85 | 27.65 |
| Saturated Fat (gm%) | 0.0558 | **8.53 | 8.74 |
| Cholesterol (gm%) | 0 | 0.012 | 1.99 |

**Supplementary Table S2.**
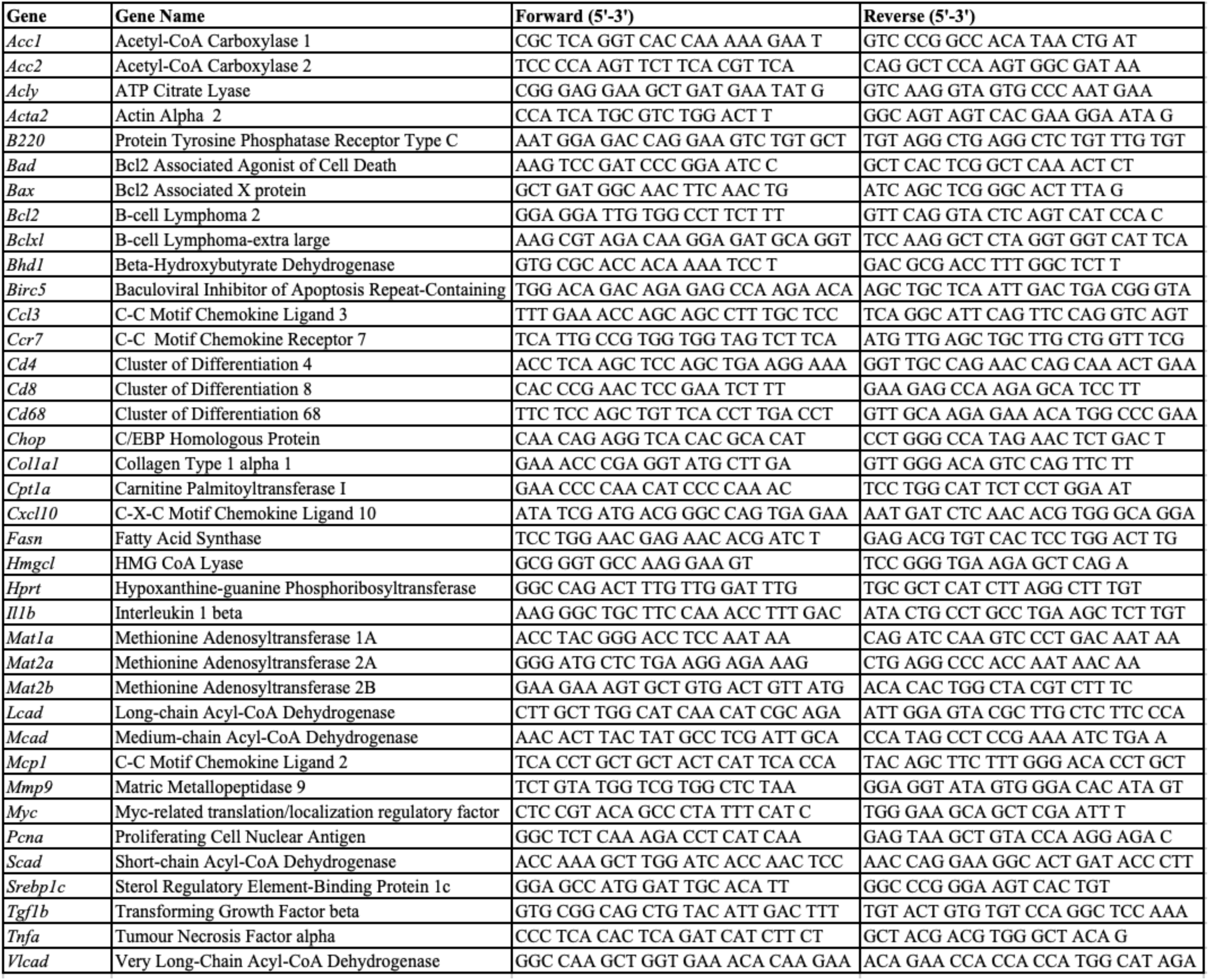
List of mouse primers used in qPCR experiments.

**Figure S1.**
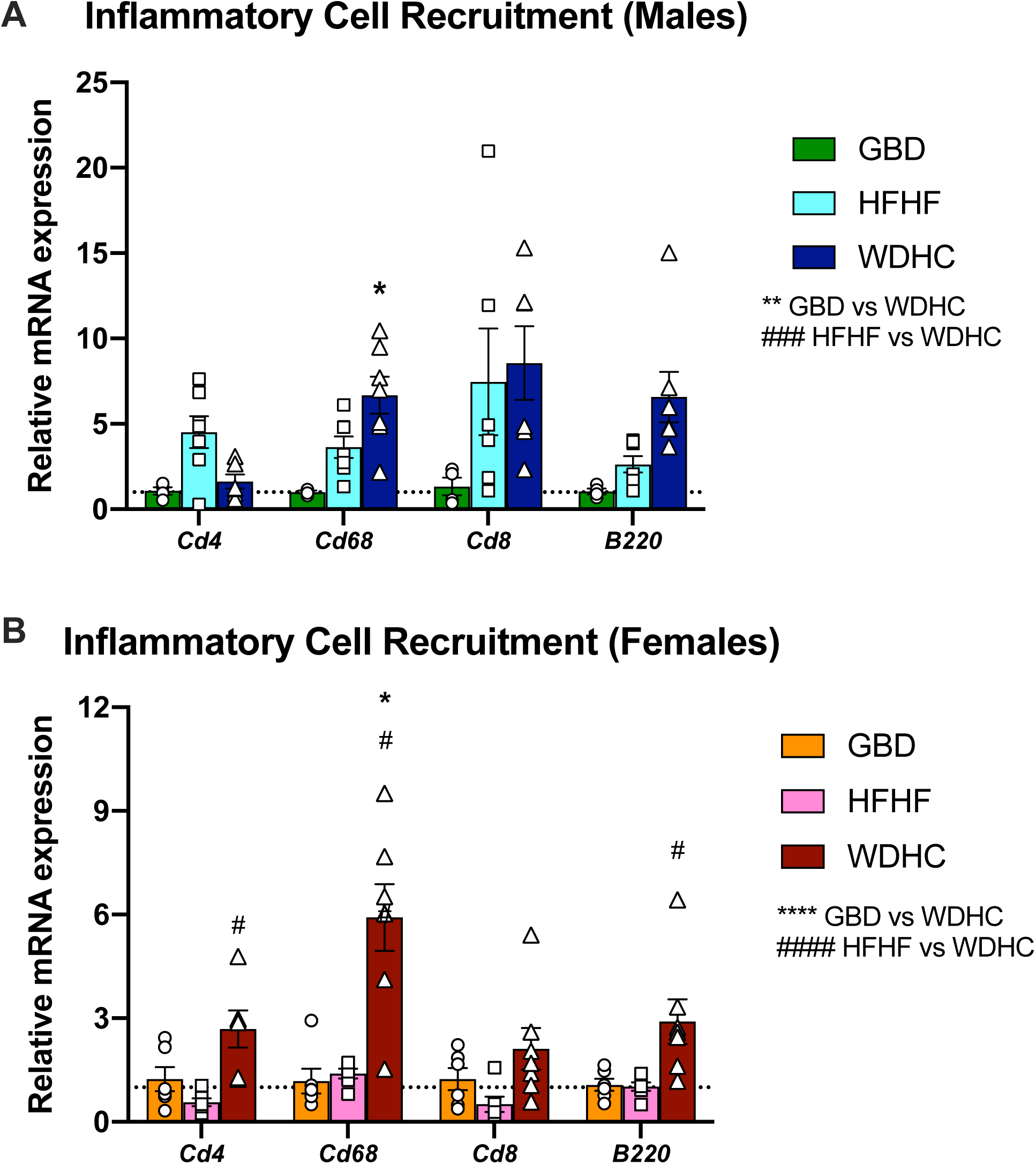
WDHC-fed mice have upregulated hepatic expression of genes involved in inflammatory cell recruitment. Gene expression measured by qPCR in (A) male and (B) female mice. Data are expressed as mean ± SEM (n=4-8) and representative of two individual cohorts. Results of Two-way ANOVA and t-tests are reported in panels. ^*^p ≤ 0.05, ^**^p ≤ 0.01, ^****^p ≤ 0.0001 compared to GBD; ^#^p ≤ 0.05, ^####^p ≤ 0.0001 comparing WDHC to HFHF.

**Figure S2.**
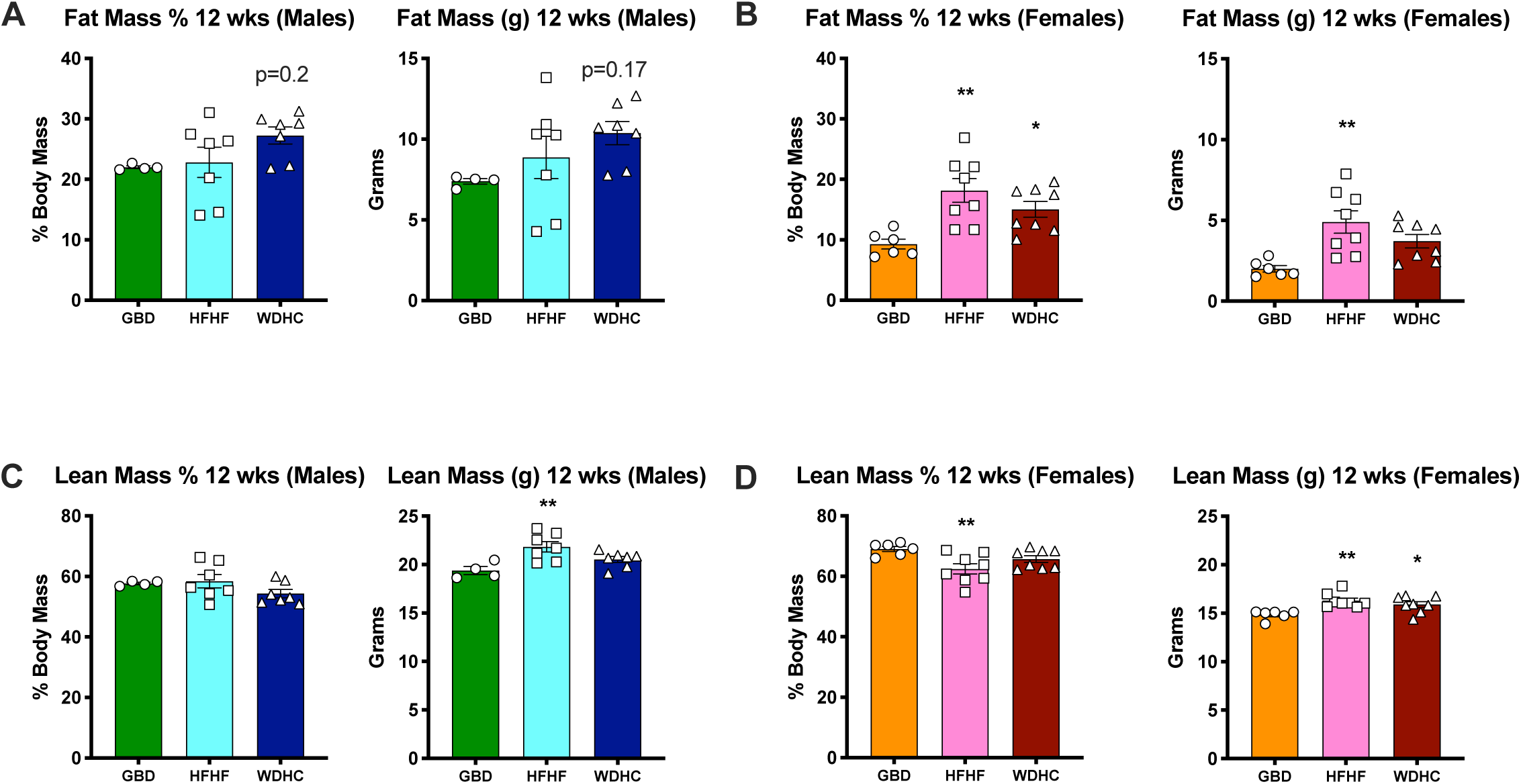
Sex-specific differences in lean and fat mass are observed after 12 weeks of diet feeding. (A-B) Fat and (C-D) lean masses (normalized to body mass and absolute values) taken at 12 weeks of feeding by MRI in male (A, C) and female mice (B, D). Data are expressed as mean ± SEM (n=4-8) and representative of two individual cohorts. Results of One-way ANOVA are reported in panels. ^*^p ≤ 0.05, ^**^p ≤ 0.01 compared to GBD.

**Figure S3.**
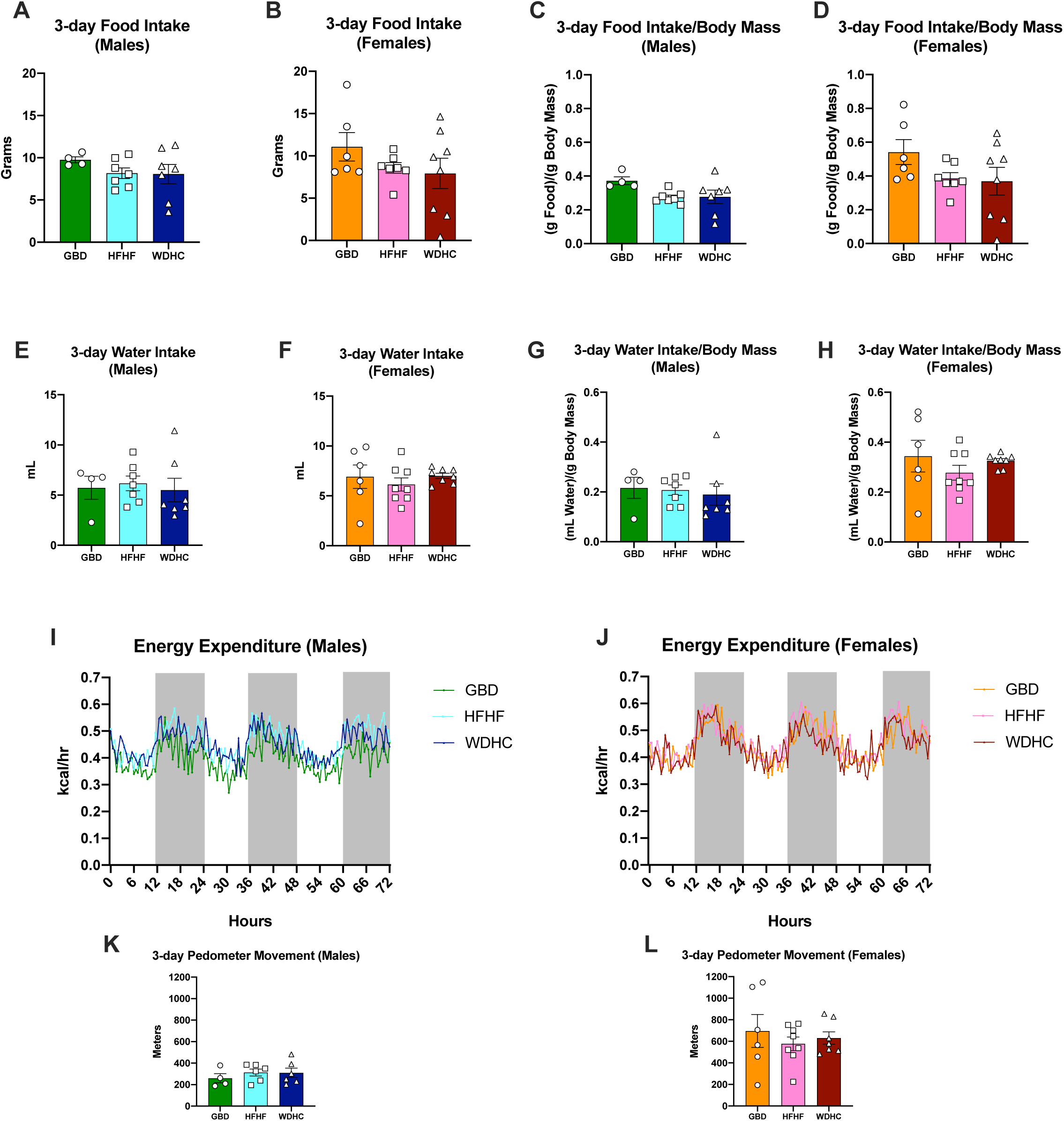
Energy intake and expenditure were not different between mice on different diets. (A-B) 3-day total food intake; (C-D) 3-day total food intake normalized to body mass; (E-F) 3-day total water intake; (G-H) 3-day total water intake normalized to body mass; (I-J) Energy expenditure over 3 days; and (K-L) 3-day total pedometer movement were measured using metabolic cages. Data are expressed as mean ± SEM (n=4-8) and representative of two individual cohorts.

**Figure S4.**
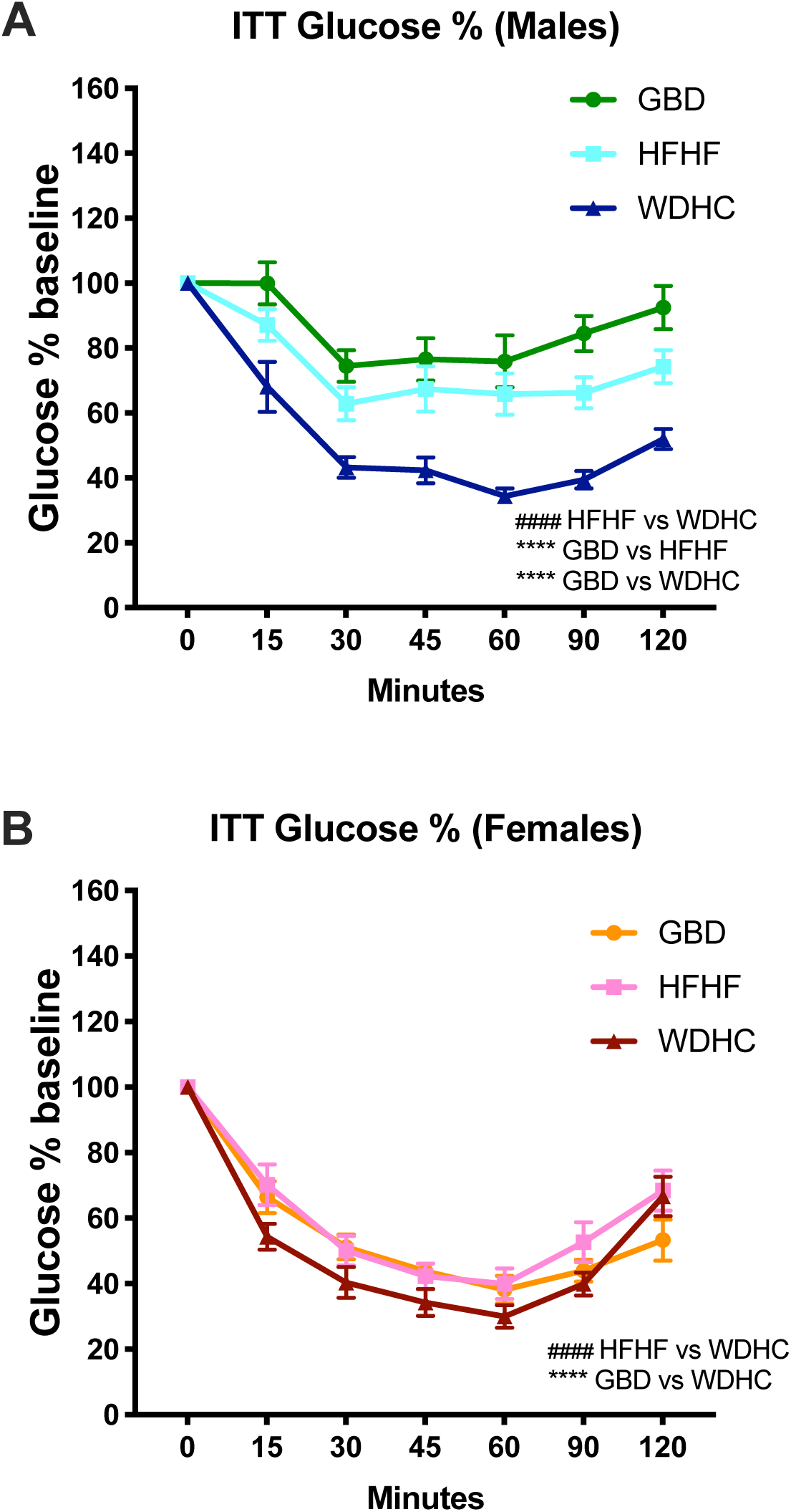
WDHC-fed mice were more insulin sensitive than GBD- and HFHF-fed mice. Insulin Tolerance Tests (ITTs) for (A) males and (B) females presented as glucose levels normalized to baseline values (5hr fast). Data are expressed as mean ± SEM (n=4-8) and representative of two individual cohorts. Results of Two-way ANOVA are reported in panels. ^****^p ≤ 0.0001 compared to GBD; ^#^p ≤ 0.05, ^####^p ≤ 0.0001 comparing WDHC to HFHF.

**Figure S5.**
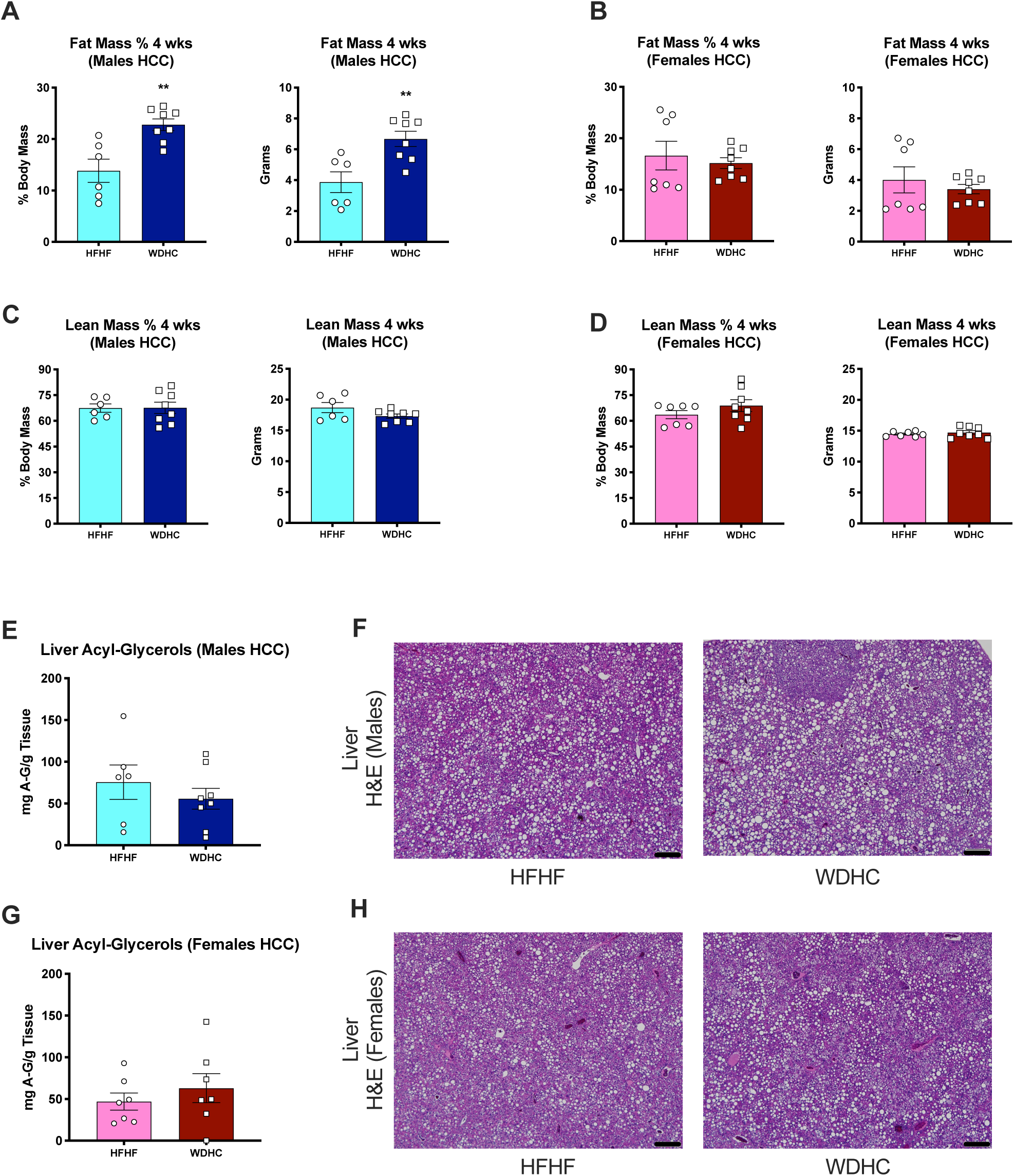
WDHC-fed and HFHF-fed mice with HCC had similar levels of hepatic acyl-glycerols. (A-B) Whole body fat mass percentage and absolute fat mass by MRI after 4 weeks on the diets; (C-D) lean mass percentage and absolute lean mass taken with MRI at 4 weeks on the diets. (E) Male liver acyl-glycerol content at 24 weeks of feeding and (F) H&E staining of male liver tissue at 24 weeks of feeding (bars represent 200um). (G) Female liver acyl-glycerol content at 24 weeks of feeding, and (H) H&E staining of female liver tissue at 24 weeks of feeding. Data are expressed as mean ± SEM (n=6-8) and representative of two individual cohorts. Results of t-tests are reported in panels. ^**^p ≤0.01, ^***^p ≤0.001.

**Figure S6.**
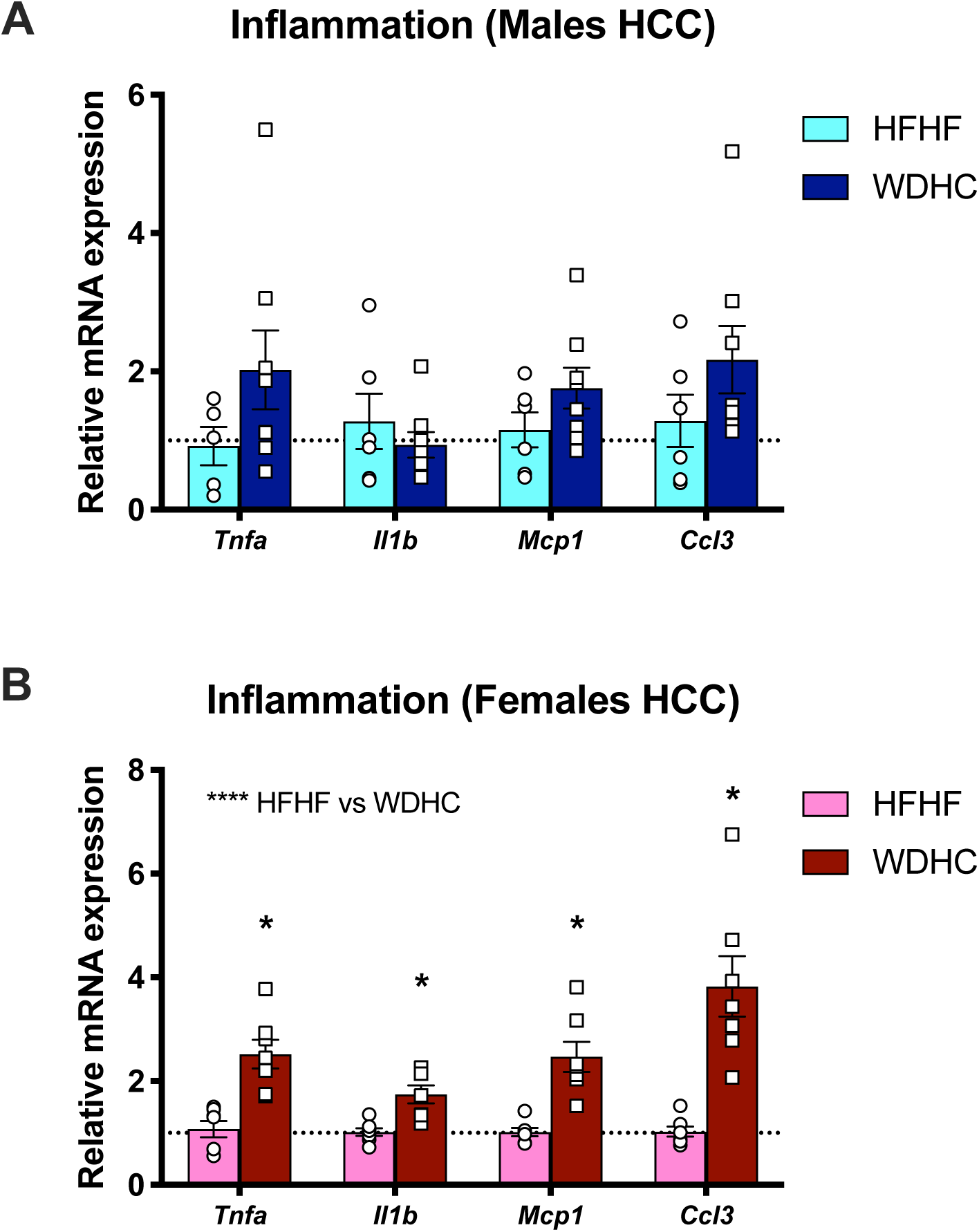
WDHC-fed females have increased expression of inflammatory genes in their livers compared to HFHF-fed mice. Hepatic expression of inflammatory genes measured by qPCR in (A) male and (B) female mice. Data are expressed as mean ± SEM (n=4-8) and representative of two individual cohorts. Results of Two-way ANOVA and t-tests are presented in panels. ^*^p ≤ 0.05, ^****^p ≤ 0.0001.

